# Transfer of cyclobutane pyrimidine dimer photolyase to chloroplasts for *Poaceae* survival under ultraviolet-B radiation

**DOI:** 10.1101/2023.09.18.558314

**Authors:** Momo Otake, Mika Teranishi, Chiharu Komatsu, Mamoru Hara, Kaoru Okamoto Yoshiyama, Jun Hidema

## Abstract

Cyclobutane pyrimidine dimer (CPD) photolyase (PHR), the primary enzyme for repairing the CPD induced by ultraviolet B (UV-B) radiation, is essential for plants living under sunlight. Rice CPD photolyase (OsPHR), is such a unique triple-targeting protein. The signal sequences required for its translocation to the nucleus or mitochondria are located in the C-terminal region but were yet to be identified for chloroplasts. Here, we identified sequences located in the N-terminal region, including the serine-phosphorylation site at position 7 of OsPHR, and found that OsPHR is transported/localized to chloroplasts via a vesicle transport system under the control of serine phosphorylation. However, the sequence identified in this study is only conserved in some *Poaceae* species and in many other plants, PHR does not localize to chloroplasts Therefore, we reasoned that *Poaceae* species need the ability to repair CPD in the chloroplast genome to survive under sunlight and have acquired this new mechanism for chloroplast translocation.

## Introduction

Plants use sunlight as an energy source for photosynthesis and are inevitably exposed to harmful ultraviolet B (UV-B) radiation (280-315 nm). Although UV-B is a minor fraction of the UV radiation that reaches the Earth’s surface, it causes substantial damage to plant macromolecules such as lipids, proteins, and membranes, especially DNA (Teremura, 1983). The cyclobutane pyrimidine dimer (CPD) represents a major type of DNA damage induced by UV-B radiation (Britt, 1996), which induces CPD on DNA, and hence in all organelles containing DNA (Stapleton et al., 1994; Takahashi et al., 2011). The accumulation of DNA damage induces mutations or cell death by impeding replication and transcription, thereby inhibiting plant growth and development. Cyclobutane pyrimidine dimers (CPDs) are the principal cause of UV-B-induced growth inhibition in rice plants grown under supplementary UV-B radiation (Hidema et al., 2007; Teranishi et al., 2012; Mmbando et al., 2021).

Plants have developed mechanisms to cope with UV-B-induced CPDs, including photoreactivation and nucleotide excision repair (NER) (Britt, 1996). In higher plants, photoreactivation is the primary mechanism for repairing CPDs because the rate of photoreactivation is faster than that of NER. Photoreactivation is mediated by photolyase (PHR), which absorbs ultraviolet A (UV-A) (315-400 nm) and blue radiation as an energy to monomerize dimers (Britt, 1996; Sancar et al., 2004). PHR deficient *Arabidopsis* (Landry et al., 1997; Dündar et al., 2020) or rice (Hidema et al., 2007) plants show UV-B hypersensitive phenotype. We previously demonstrated that the difference in UV-B sensitivity among rice cultivars depends on PHR activity in the cell (Hidema et al., 2000; Teranishi et al., 2004; Mmbando et al., 2020) and that elevated PHR activity can markedly alleviate UV-B-induced growth inhibition in rice (Hidema et al., 2007; Teranishi et al., 2012; Mmbando et al., 2021). Thus, CPD photolyase is an essential protein for plants grown under sunlight.

CPD photolyase is widely distributed among species ranging from eubacteria and archaebacteria to eukaryotes, apart from eutherian mammals (Yasui et al., 1994). Eukaryotic cells have at least two organelles containing DNA: nuclei and mitochondria. In addition to the nuclear and mitochondrial genomes, photosynthetic eukaryotic cells possess chloroplasts as organelles that contain DNA. Yeast CPD photolyase is transported into each organelle via a signal sequence for the nuclei or mitochondria, and yeast efficiently photorepairs CPDs on both nuclear and mitochondrial DNA using CPD photolyase (Yasui et al., 1992). In rice plants, rice CPD photolyase (OsPHR), which is encoded by a single-copy gene and not a splice variant, is expressed and targeted not only to the nucleus and mitochondria but also to chloroplasts (Takahashi et al., 2011). This protein repaired UV-B-induced CPDs in all three genomes. In addition, we had previously identified the nuclear and mitochondrial target signal sequences retained in the C-terminal region of OsPHR and found that each OsPHR target sequence was highly conserved among plant species (Takahashi et al., 2014). However, the location and nature of the chloroplast-targeting information contained within OsPHR remained unknown.

In this study, we searched for sequences required for OsPHR to be transferred to chloroplasts using a systematic deletion analysis. We found that the sequence is located within the N-terminal region and that the 7th serine residue and neighboring prolines of OsPHR are important for its translocation through the endoplasmic reticulum (ER) to the chloroplast. In addition, the phosphorylation of serine at position 7 of OsPHR (Takahashi et al., 2008; Teranishi et al., 2013) would be involved in the regulation of OsPHR chloroplast translocation via the ER-Golgi system. However, the amino acid sequence of the N-terminal region of the PHR varies widely among plant species, and the sequences identified in this study were not highly conserved among plant species. We examined the chloroplast localization of PHR in various plants using transient expression analysis and found that PHR was not localized to the chloroplasts in many plants, except for *Poaceae*. It is thought that *Poaceae* have acquired a mechanism to protect chloroplasts from UV-B-induced damage by transferring PHR to chloroplasts on their own during evolution.

## Results

### N-terminal region sequence is required for chloroplast translocation of OsPHR

We have previously reported the possibility that the sequence required for OsPHR transfer to the chloroplast is located in the N-terminal region of OsPHR (Takahashi et al., 2014). Using TargetP (https://services.healthtech.dtu.dk/services/TargetP-2.0/) (Armenteros et al., 2019), a popular protein localization prediction algorithm, we analyzed the presence of sequences predicted to be transferred to the chloroplasts within the N-terminal region of OsPHR (amino acids 1-47). It was deduced that the sequence within amino acids 1-47 of OsPHR is most likely transported to the ER, but not to the chloroplasts. Thus, we first focused on N-terminal region of OsPHR and generated the transgenic rice plants stably expressing citrine fused to the carboxy (C) terminus of a fragment of the N-terminus OsPHR (amino acids 1-47, N_1-47_-OsPHR) under the cauliflower mosaic virus (CaMV) promoter to determine whether the sequence of the N-terminal region of OsPHR can be transferred to chloroplasts. In transgenic rice stably expressing citrine fused to the C-terminus of full-length (FL) OsPHR under the CaMV promoter (FL-OsPHR), the citrine fluorescence of FL-OsPHR was clearly detected in the chloroplasts of rice leaf cells (Fig. 1A). Compared with the localization of citrine using a control vector, the citrine fluorescence of N_1-47_OsPHR was also clearly detected in the chloroplasts of rice leaf cells (Fig. 1A). Therefore, to examine whether OsPHR is translocated to chloroplasts using the sequence of the N-terminal region, transgenic rice plants expressing citrine fused to the C-terminus of partial OsPHR without amino acids 1-14 (Δ_1-14_OsPHR) were generated. When citrine fluorescence of Δ_1-14_OsPHR was observed in rice cells, citrine fluorescence was not detected in the chloroplasts, whereas it was detected in the nuclei and mitochondria (Fig. 1B). Furthermore, when the UV-B sensitivity of the transgenic lines expressing Δ_1-14_OsPHR (lines 11 and 13) was compared with that of plants expressing FL-OsPHR, obvious leaf browning was observed in the transgenic lines expressing Δ_1-14_OsPHR. Both transgenic rice lines showed a UV-B-sensitive phenotype compared with the transgenic line with chloroplast-translocated OsPHR (FL-OsPHR) (Fig. 1C). Therefore, it is strongly suggested that the sequence required for OsPHR to translocate to chloroplasts in rice cells is located within the N-terminal region and that the translocation of OsPHR to chloroplasts is important for UV-B sensitivity in rice.

**Figure 1.**
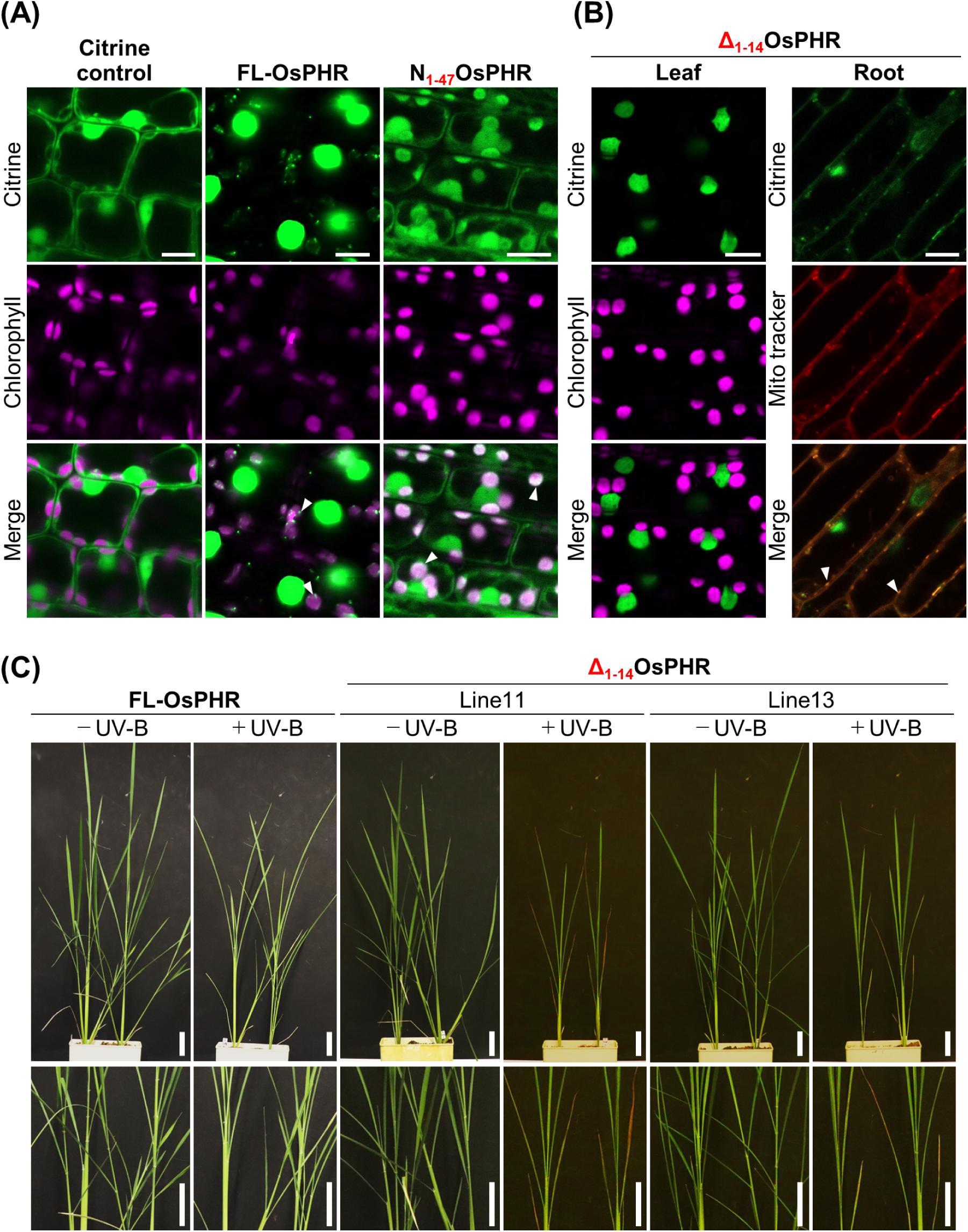
The N-terminal 1-14 amino acid sequences are critical for chloroplast translocation of OsPHR and affects UV-B sensitivity. (A) and (B) show the cellular localization of OsPHR. Conforcal microscopic images of the transgenic rice plants expressing citrine fused to the C terminus of full length (FL) OsPHR (amino acids 1-506, FL-OsPHR) and a fragment of OsPHR (N_1-47_ or Δ_1-14_ OsPHR) in each case under the control of a CaMV 35S promoter. Arrowheads indicate the signal of Citrine in chloroplast (a) and mitochondria (b). Bars = 10 μm. (C) Effects of UV-B radiation on rice strains (FL-OsPHR, and lines 11 and 13 of Δ_1-14_ OsPHR). The rice plants were grown in the growth cabinet for 30 d with (+UV-B: 1.0 W m^-2^) or without (-UV-B) supplementary UV-B radiation. Bars = 5 cm.

### Chloroplast translocation of OsPHR is inhibited by brefeldin A (BFA) treatment

As mentioned previously, the N-terminal region of OsPHR contains sequences that are likely to be transported to the ER. Recently, it has been reported that a part of the nuclear-encoded chloroplast protein is imported from the ER-Golgi system to the chloroplast through the secretory pathway (Kleffmann et al., 2004; Friso et al., 2004; Asatsuma et al., 2005) or is incorporated into chloroplasts without cleavage of the peptide within the N-terminal region of the protein (Armbruster et al., 2009). Therefore, to deduce the transport pathway of OsPHR to chloroplasts, we analyzed the effects of BFA, a fungal antibiotic that inhibits Golgi-mediated vesicular traffic (Ritzenthaler et al., 2002), on the translocation of OsPHR to chloroplasts. We constructed an expression vector expressing FL-OsPHR and performed a transient expression assay using protoplasts prepared from shoot of rice seedlings. When citrine alone was introduced into rice protoplasts as a control, citrine fluorescence was observed not only in the cytoplasm but also in the nucleus (Kaiser et al., 2009) (Fig. 2A). Relatively small proteins often enter the nucleus passively. In contrast, when an FL-OsPHR expression vector was introduced into rice protoplasts, citrine fluorescence was detected in the nuclei and chloroplasts as nucleoids. When the protoplasts were treated with 0.1% dimethyl sulfoxide (DMSO) as a control, citrine signals were detected in the nuclei and chloroplasts of approximately 76% of the transformed protoplasts (Fig. 2B and C). In contrast, in BFA-treated protoplasts, the number of chloroplasts in which citrine fluorescence was detected reduced to approximately 12%. These results indicate that OsPHR is translocated to chloroplasts via the ER-Golgi system through the secretory pathway.

**Figure 2.**
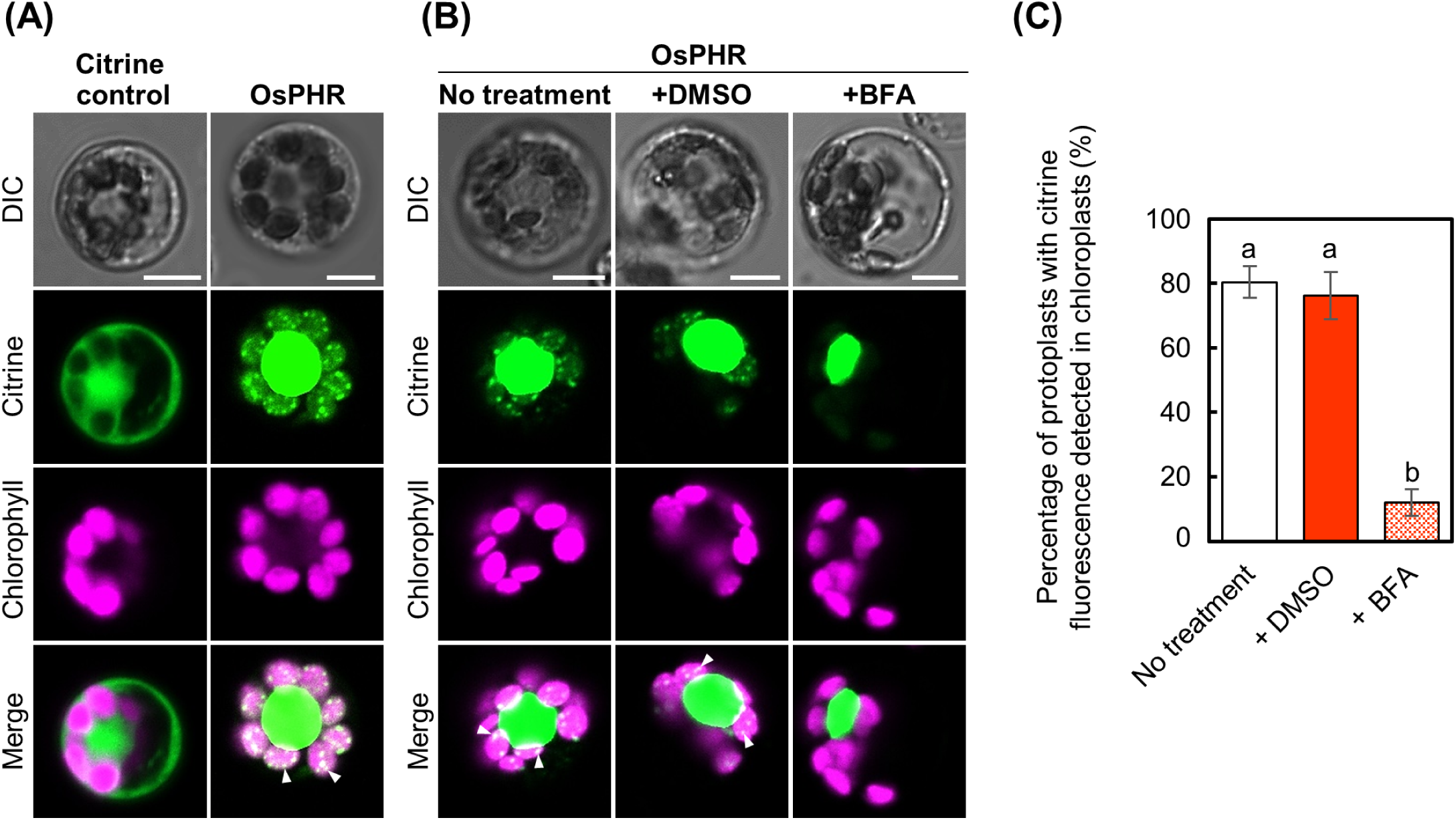
Chloroplast translocation of OsPHR is inhibited by BFA treatment. (A) and (B) Confocal microscopic images of rice protoplasts expressing citrine (control) or FL-OsPHR under the control of the CaMV 35S promoter (A), and of the protoplasts, expressing FL-OsPHR, treated with DMSO as a control or BFA. Arrowheads indicate the signal of citrine in chloroplast. Bars = 5 µm. (C) The percentage of protoplasts in which citrine fluorescence is detected in chloroplasts among all transformed protoplasts treated with DMSO and BFA (±se, n ≧80 protoplasts). Different letters in each graph denote significant differences from each other based on Tukey’s test (*P* < 0.05).

## Proline residues are important for chloroplast translocation of OsPHR

Certain proteins are transported to the mitochondria via the ER-Golgi transport system, which is characterized by the presence of a hydrophobic region (H-region) within the N-terminal region (Martoglio et al., 1998; Kida et al., 2000; Kanaji et al., 2000). In addition, proline residues preceding the H-region are critical for passage through the ER (Ritzenthaler et al., 2002). The N-terminal region of OsPHR is rich in hydrophobic amino acids, particularly proline. Hence, to investigate whether proline residues in the N-terminal region of OsPHR are involved in chloroplast translocation, a transient expression vector encoding a mutated OsPHR was constructed by replacing the proline residue with an alanine residue in the 1-14 amino acid region, which is essential for chloroplast translocation (Fig. 1B). The results showed that replacing the 13th proline with an alanine (P13A-OsPHR) did not alter the chloroplast translocation of OsPHR, whereas replacing the 2nd, 3rd, or 8th proline (P2,3A- or P8A-OsPHR) slightly inhibited chloroplast translocation (Fig. 3A-C). In contrast, when the 9th proline was converted to alanine (P9A-OsPHR), the chloroplast translocation of OsPHR was inhibited to a greater extent than when proline at other positions was converted to alanine, and the degree of inhibition increased further when both the 8th and 9th proline residues were converted to alanine (P8,9A-OsPHR) (Fig. 3A-C). These results suggested that the 8th and 9th prolines of OsPHR play notable roles in the translocation of OsPHR to the chloroplasts.

**Figure 3.**
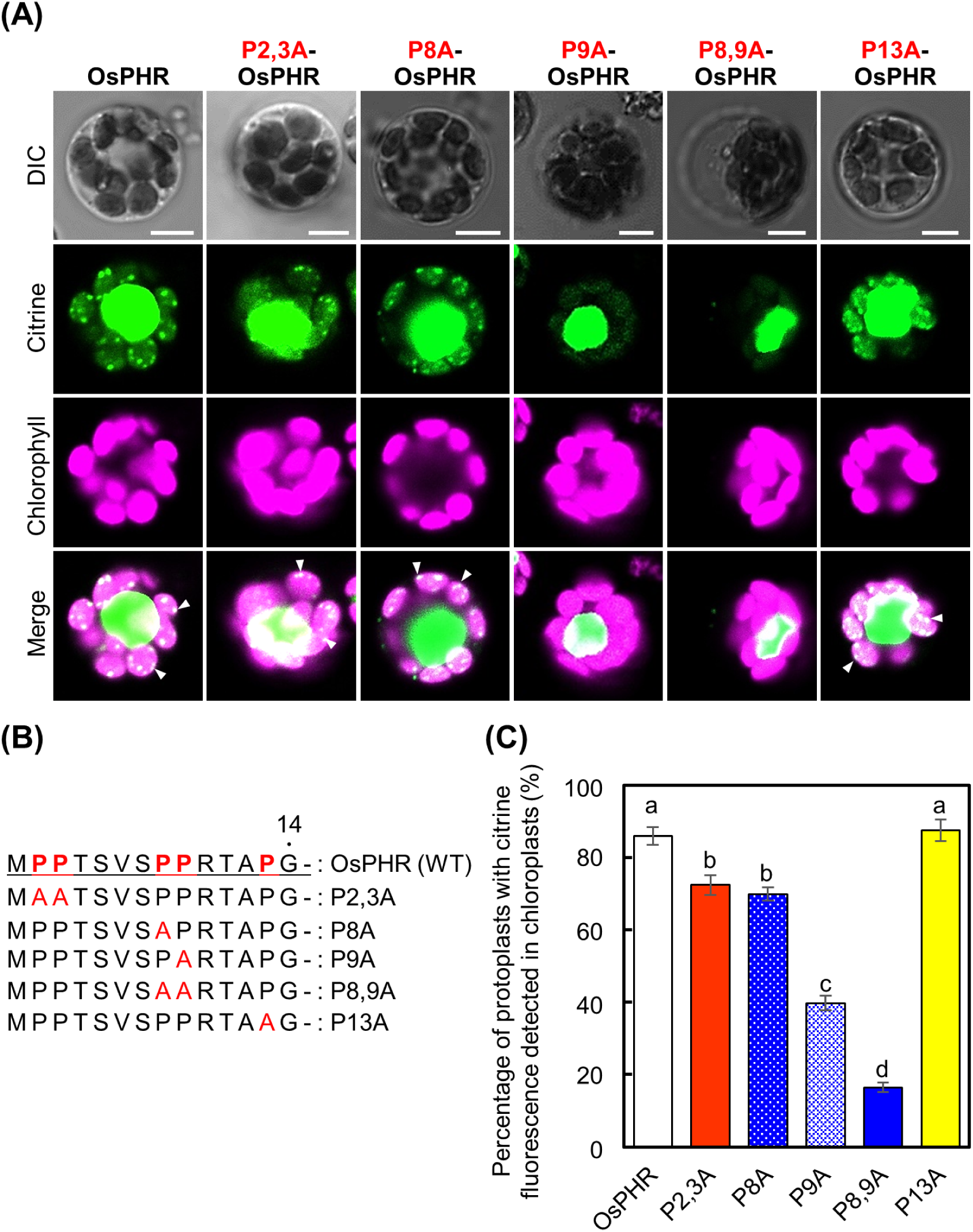
Proline residues are important for chloroplast translocation of OsPHR. (A) Confocal microscopic images of rice protoplasts expressing FL-OsPHR and a mutated FL-OsPHR by replacing a proline residue with an alanine residue (P2,3A, P8A, P9A, P8,9A and P13A) under the control of the CaMV 35S promoter. Arrowheads indicate the signal of citrine in chloroplast nucleoid. Bars = 5 μm. (B) Proline residues were mutated into alanine residues in 1-14 amino acid region of OsPHR as indicated. (C) The percentage of protoplasts in which citrin fluorescence of OsPHR or OsPHR by replacing a proline residue with an alanine residue (P2,3A, P8A, P9A, P8,9A or P13A) is detected in chloroplasts among all transformed protoplasts (±se, n ≧140 protoplasts). Different letters in each graph denote significant differences from each other based on Tukey’s test (P < 0.05).

### Phosphorylated OsPHR suppresses chloroplast translocation

We have previously shown that phosphorylated and non-phosphorylated forms of the 7th serine of OsPHR are present in vivo (Takahashi et al., 2008; Teranishi et al., 2013) and that there is a difference in the quantity of phosphorylated OsPHR in the chloroplasts, mitochondria, and nuclei of rice cells (Takahashi et al., 2011). However, when, where, and by what kinase OsPHR is phosphorylated and the biochemical and physiological functions of phosphorylated PHR are unclear. These results raise the possibility that the phosphorylation state of CPD photolyase might be associated with its targeting to different organelles. Therefore, to confirm whether the difference in the phosphorylation state of the 7th serine of OsPHR affected the chloroplast translocation of OsPHR, an expression vector expressing FL-OsPHR with the 7th serine replaced by alanine (pseudo-unphosphorylated state: S7A-OsPHR), aspartic acid (pseudo-phosphorylated state: S7D-OsPHR), or glutamic acid (pseudo-phosphorylated state: S7E-OsPHR) under the control of the CaMV promoter was introduced into rice protoplasts. When pseudo-unphosphorylated S7A-OsPHR was introduced into rice protoplasts, the percentage of protoplasts in which citrine fluorescence was detected in the chloroplasts was approximately the same as that in unmutated OsPHR (Fig. 4A-C). In contrast, when pseudo-phosphorylated S7D-OsPHR or S7E-OsPHR was introduced into rice protoplasts, the percentage of protoplasts in which citrine fluorescence was detected in the chloroplasts was dramatically reduced compared to that of OsPHR or pseudo-unphosphorylated S7A-OsPHR (Fig. 4A-C). Furthermore, when expression vectors expressing S7A-OsPHR were introduced into rice protoplasts and treated with DMSO or BFA, the percentage of protoplasts in which the fluorescent signal of S7A-OsPHR was detected in the chloroplasts was reduced by BFA treatment (Fig. 4D and E). These results indicate that OsPHR with an unphosphorylated serine at position 7 is more clearly translocated to the chloroplasts via the ER-Golgi system than phosphorylated OsPHR. In addition, to confirm whether OsPHR in the unphosphorylated state of the 7th serine is more actively translocated to chloroplasts in vivo than phosphorylated OsPHR, we generated transgenic rice plants stably expressing S7A-OsPHR or S7D-OsPHR under the CaMV promoter. In transgenic rice plants introduced with S7A-OsPHR, lines 7 and 9, a strong signal of citrine fluorescence was clearly detected in the chloroplasts and nuclei of rice cells, similar to the OsPHR-transgenic rice (Fig. 4F). In contrast, in transgenic rice plants transformed with S7D-OsPHR (lines 3 and 4), citrine fluorescence was detected in the nucleus but not in the chloroplasts (Fig. 4F). Thus, the phosphorylation of serine at position 7 of OsPHR is involved in the regulation of OsPHR chloroplast translocation.

**Figure 4.**
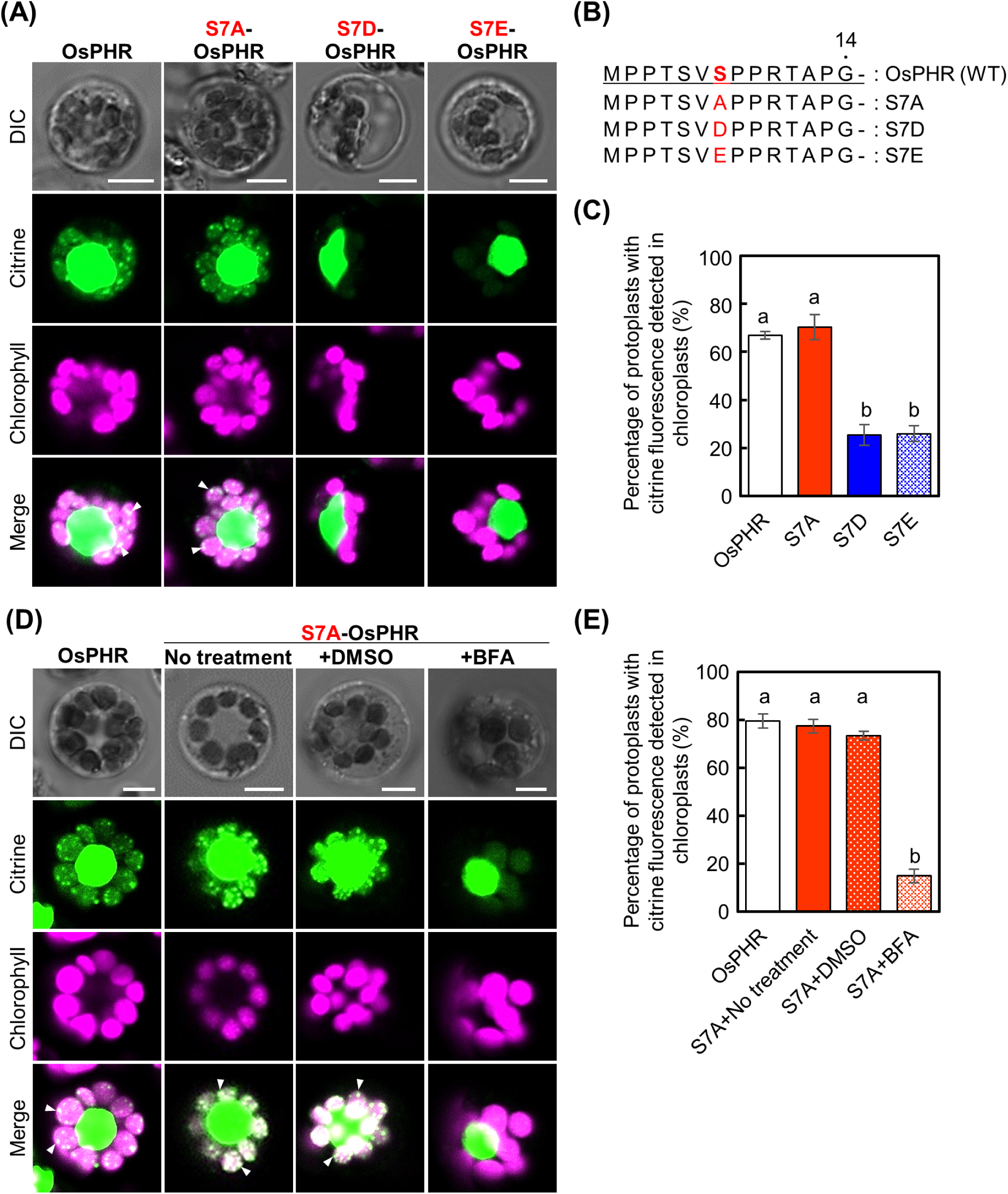
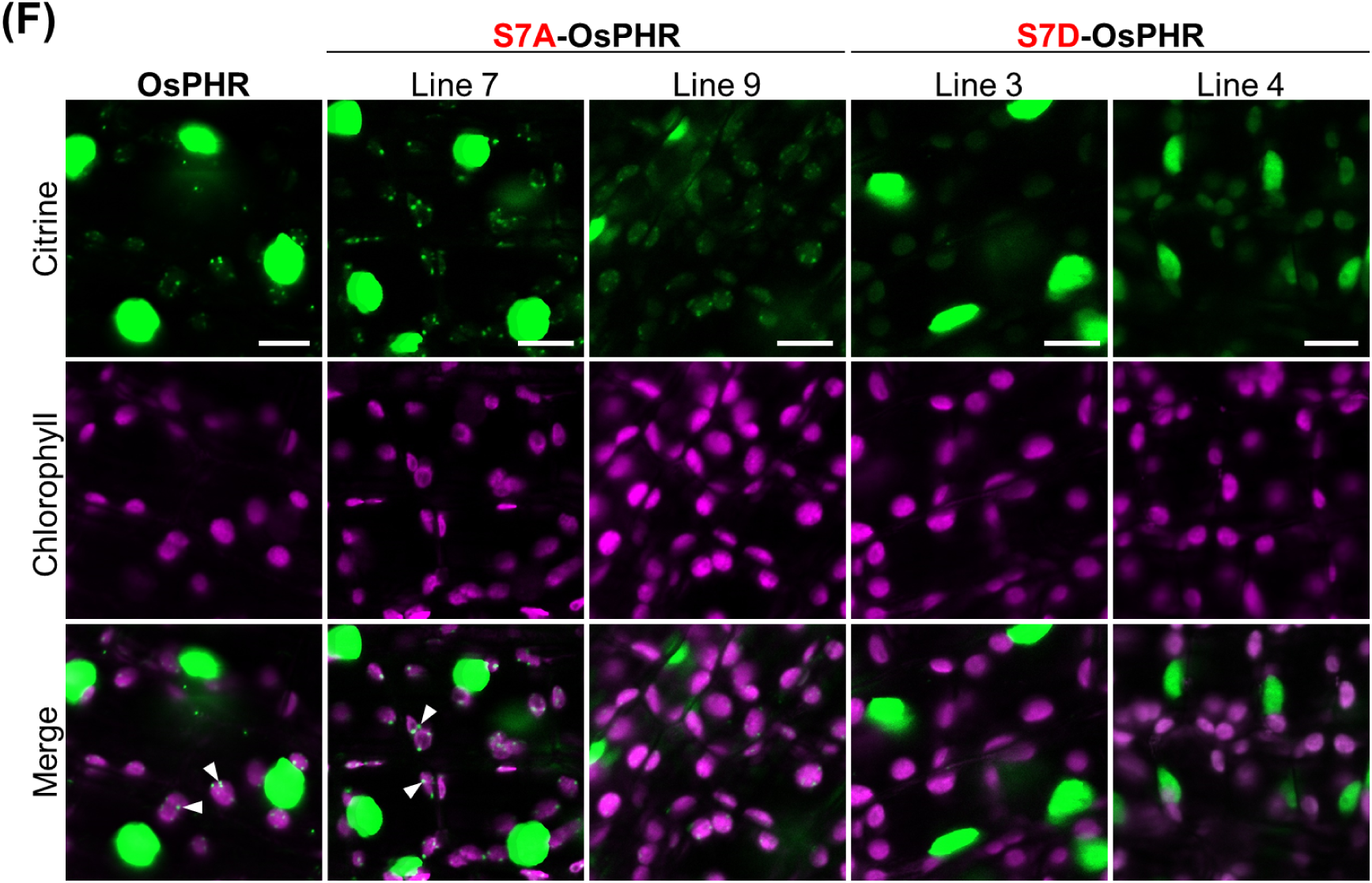
Phosphorylation state of the 7th serine is involved in the chloroplast translocation of OsPHR. (A) Conforcal microscopic images of rice protoplasts expressing OsPHR-Citrine, pseudo-unphosphorylated S7A-OsPHR-Citrine, pseudo-phosphorylated S7D- or S7E-OsPHR-Citrine in each case under the control of the CaMV 35S promoter. Bars = 5 μm. (B) The phosphorylation state of the seventh serine residues was mutated into alanine, aspartic acid, and glutamic acid residues of OsPHR as indicated. (C) The percentage of protoplasts in which citrine fluorescence of OsPHR, S7A-OsPHR, S7D-OsPHR or S7E-OsPHR is detected in chloroplasts among all transformed protoplasts (±se, ≧150 protoplasts). (D) Conforcal microscopic images of rice protoplasts are treated with DMSO or BFA expressing S7A-OsPHR-Citrine. Bars = 5 μm. (E) The percentage of protoplasts in which citrine fluorescence of S7A-OsPHR are detected in chloroplasts among all transformed protoplasts treated with DMSO and BFA (±se, ≧200 protoplasts). (F) Conforcal microscopic images of the transgenic rice plants expressing citrine fused to the C terminus of OsPHR, S7A-OsPHR and S7D-OsPHR under the control of CaMV 35S promoter. Bars = 10 μm.

### Chloroplast translocation of PHR depends among plant species

OsPHR was shown to translocate to chloroplasts using the sequence of the N-terminal region. However, the amino acid sequence of the N-terminal region of PHR varied widely among plant species, as shown in Fig. 5A and Supplemental Fig. S1. Interestingly, only plants belonging to the *Poaceae* family, such as wheat and barley, have PHRs that show high homology with the amino acid sequence of the N-terminal region, which is important for the transfer of OsPHRs to chloroplasts. On the other hand, the sequences of amino acids 1-14 of OsPHR are missing from the *Arabidopsis* PHR (AtPHR). This implies that most plant PHR are not translocated to chloroplasts or that the sequence required for OsPHR to transfer to chloroplasts may differ among plant species. Therefore, we investigated chloroplast translocation of PHR in various plants. We first constructed an expression vector which expressed citrine fused to the C-terminus of AtPHR or *Lotus japonicus* PHR (LjPHR) and performed a transient expression assay using rice protoplasts (Fig. 5B). Citrine fluorescence of OsPHR was detected in the nuclei and chloroplasts of protoplasts. In particular, strong citrine signals in chloroplasts were detected as nucleoids. In contrast, citrine fluorescence of AtPHR or LjPHR was not detected in the chloroplasts. The failure to detect citrine fluorescence in AtPHR or LjPHR in chloroplasts may have resulted from its transient introduction into rice protoplasts rather than into the host protoplasts. The AtPHR vector was transiently introduced into protoplasts prepared from *Arabidopsis* leaves (Fig. 5C). However, no citrine fluorescence was detected in the chloroplasts. In addition, we generated transgenic *Arabidopsis* plants stably expressing AtPHR to confirm whether AtPHR was translocated to the chloroplasts in *Arabidopsis* cells (Fig. 5D and E). Compared to the localization of citrine using a control vector, the citrine fluorescence of AtPHR was not detected in the chloroplasts (Fig. 5D), whereas fluorescence was detected in the nuclei of leaf cells and in the mitochondria of root cells (Fig. 5E). Furthermore, after irradiating *Arabidopsis* leaves of WT (Landsberg *erecta*) with UV radiation to induce CPD in chloroplast DNA, the leaves were irradiated with blue light, and the repair of CPD by blue light was analyzed by Southern blotting using T4 endonuclease (Chen et al., 1996; Hanawalt et al., 1989; Stapleton et al., 1997). The CPDs on chloroplast DNA fragments in *Arabidopsis* leaves were not removed after 24 h of blue light exposure, although the CPDs on nuclear DNA fragments were repaired (Fig. 5F and Supplemental Fig. S2). These results strongly indicate that in *Arabidopsis*, AtPHR cannot be translocated to the chloroplasts and is not repaired by CPDs in the chloroplast DNA.

**Figure 5.**
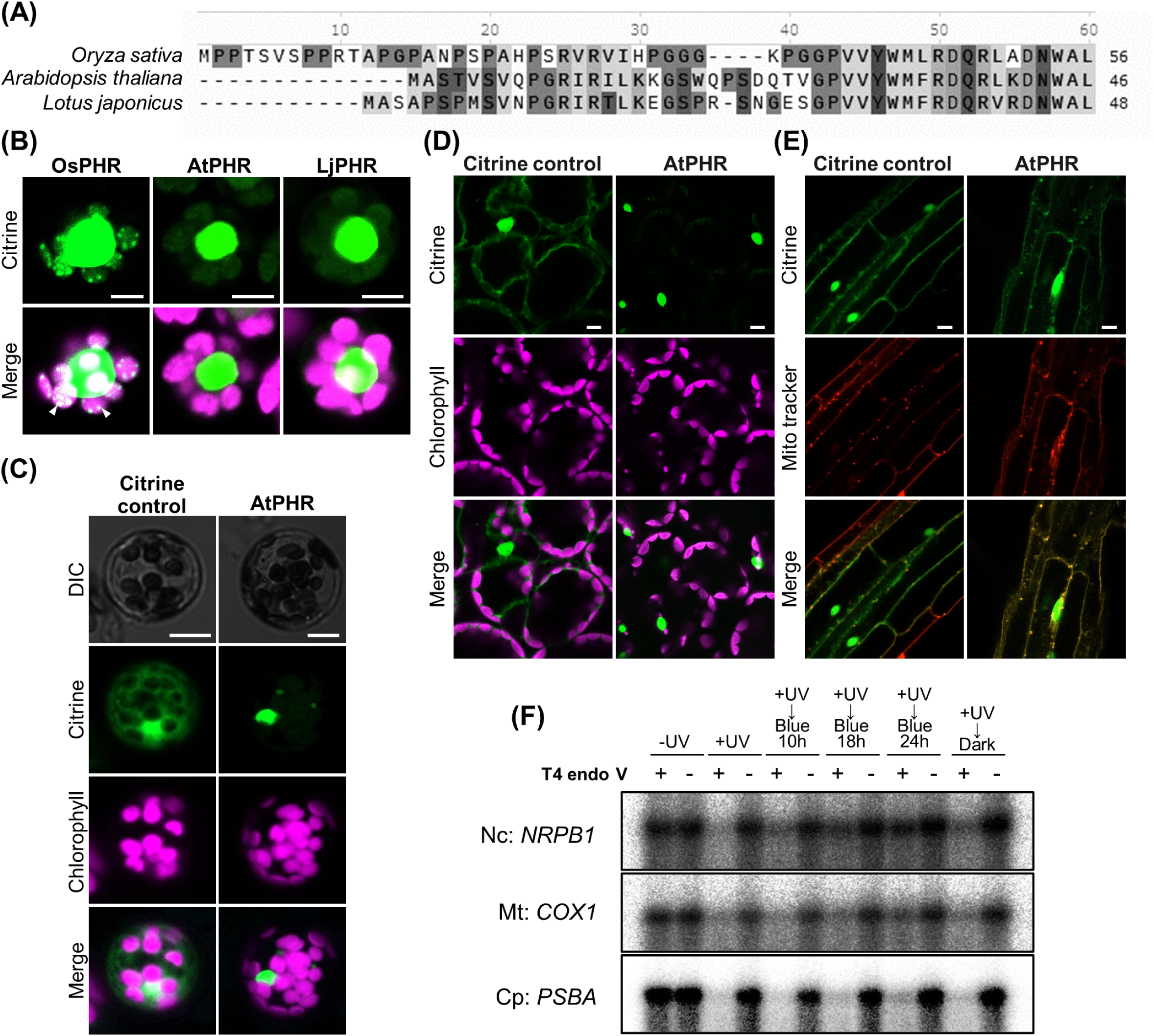
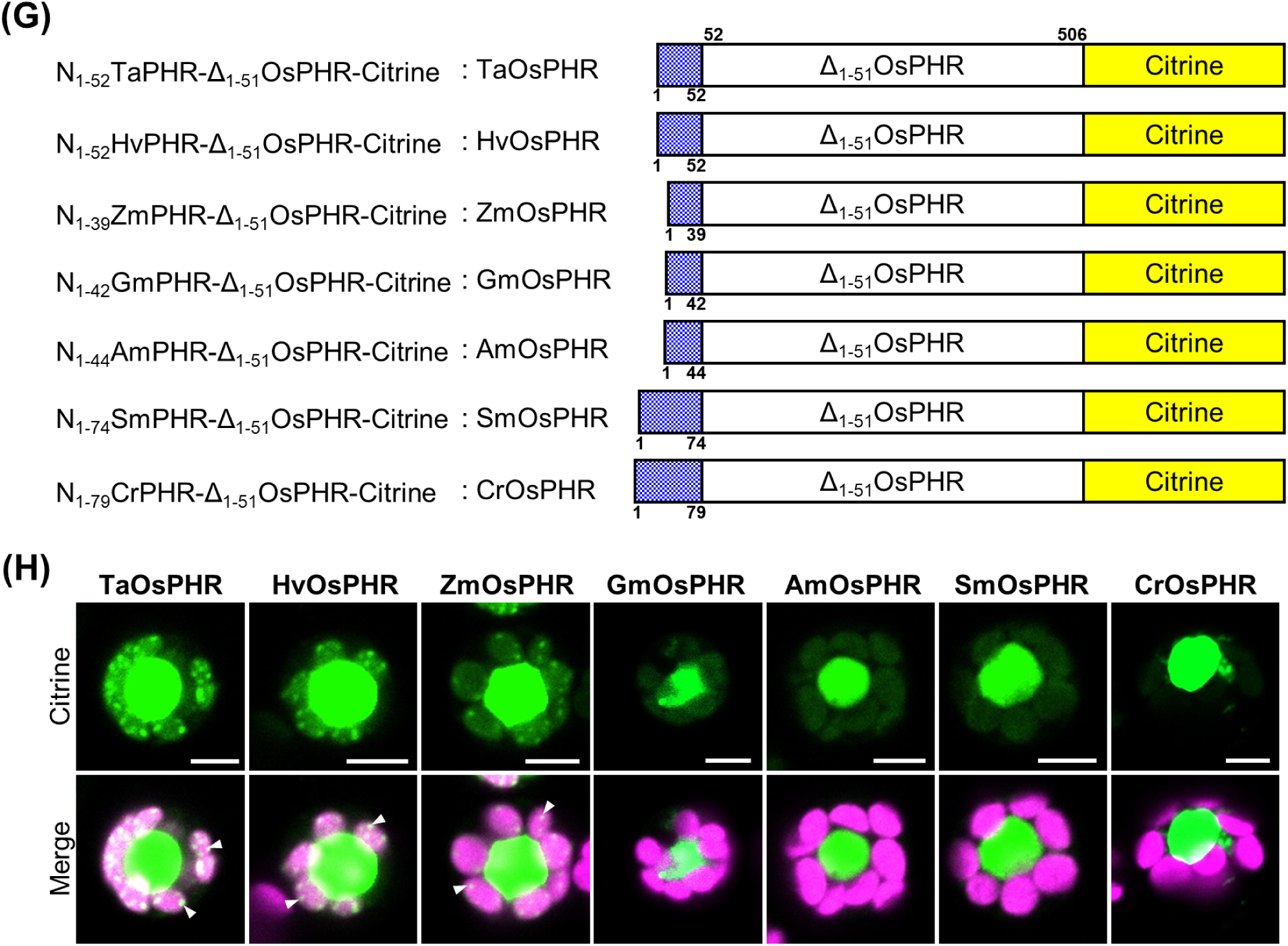
Chloroplast translocation of PHR varies among plant species. (A) Alignments of the deduced amino acid sequences at the N-terminal from *O. sativa*, *A. thaliana* and *L. japonicus*. (B) Conforcal microscopic images of rice protoplasts expressing OsPHR-Citrine, AtPHR-Citrine, and LjPHR-Citrine. Bars = 5 µm. (C) Conforcal microscopic images of *Arabidopsis* protoplasts expressing Citrine, AtPHR-Citrine, and OsPHR-Citrine. Bars = 5 µm. (D, E) Conforcal microscopic images in leaf (D) and root cells (E) of the transgenic *Arabidopsis* plants expressing Citrine fused to the C terminus of FL-AtPHR in each case under the control of CaMV 35S promoter. Bars = 10 µm. (F) Repair of CPD in *Arabidopsis* genomic DNA. Detached *Arabidopsis* leaves are harvested (no UV-C (-UV)) or are exposed to UV-C radiation (+UV) for 15 min. Leaves are then harvested immediately and exposed to blue light for 10, 18, and 24 h or kept in a light-tight box for 24 h. Genomic DNA is isolated and digested with restriction enzymes, and then the digested DNA is incubated with (+) or without (−) T4 endo V. The membranes are incubated with ^32^P-labeled probes for the nuclear-encoded genes *NRPB1*, the mitochondrial-encoded genes *COX1*, and the chloroplast-encoded gene *PSBA*. (G) Schematic drawing of the constructs used for the analysis. (H) Conforcal microscopic images of rice protoplasts expressing TaOsPHR-Citrine, HvOsPHR-Citrine, ZmOsPHR-Citrine, GmOsPHR-Citrine, AmOsPHR-Citrine, SmOsPHR-Citrine, and CrOsPHR-Citrine. Bars = 5 µm.

We further examined chloroplast localization in various plants, *Poaceae, Fabaceae, Amborellaceae, Selaginella,* and *Chlamydomonadaceae,* to understand plant species differences in the chloroplast localization of PHR. To conduct this experiment, the DNA sequences encoding the N-terminal fragments of PHR from various plants were synthesized: amino acids 1-52 from *Triticum aestivum* PHR (N_1-52_TaPHR), 1-52 from *Hordeum vulgare* PHR (N_1-52_HvPHR) and 1-39 from *Zea mays* PHR (N_1-39_ZmPHR) belonging to *Poaceae*, 1-42 from *Glycine max* PHR (N_1-42_GmPHR) belonging to *Fabaceae*, 1-44 from *Amborellaceae* PHR (N_1-44_AmPHR) belonging to *Amborellaceae*, 1-74 from *Selaginella moellendorffii Hieron* PHR (N_1-74_SmPHR) belonging to *Selaginella*, and 1-79 from *Chlamydomonas reinhardtii* PHR (N_1-79_CrPHR) belonging to *Chlamydomonadaceae*. The amino acid sequence after the 52nd amino acid sequence of OsPHR was highly homologous among plant species (Supplemental Fig. S1). Therefore, expression vectors encoding the protein chimeras of each N-terminal fragment prepared from various plants were fused to the N-terminus of OsPHR, in which amino acids 1-51 were deleted (Δ_1-51_OsPHR-Citrine). The constructs used in this analysis are as follows: N_1-52_TaPHR-Δ_1-51_OsPHR-Citrine (TaOsPHR), N_1-52_HvPHR-Δ_1-51_OsPHR-Citrine (HvOsPHR), N_1-39_ZmPHR-Δ_1-51_OsPHR-Citrine (ZmOsPHR), N_1-42_GmPHR-Δ_1-51_OsPHR-Citrine (GmOsPHR), N_1-44_AmPHR-Δ_1-51_OsPHR-Citrine (AmOsPHR), N_1-74_SmPHR-Δ_1-51_OsPHR-Citrine (SmOsPHR), or N_1-79_CrPHR-Δ_1-51_OsPHR-Citrine (CrOsPHR) (Fig. 5G). These expression constructs were transformed into rice protoplasts. In protoplasts transformed with TaOsPHR, HvOsPHR, or ZmOsPHR, citrine fluorescence was detected in the nucleoids and chloroplasts (Fig. 5H). In contrast, the citrine fluorescence of GmOsPHR, AmOsPHR, SmOsPHR, and CrOsPHR was not detected in the chloroplasts (Fig. 5H). We also examined whether BFA treatment inhibited the detection of citrine fluorescence in plants in which citrine fluorescence was detected in chloroplasts, such as TaOsPHR, HvOsPHR, and ZmOsPHR. In these *Poaceae* species, the percentage of protoplasts in which citrine fluorescence was detected in the chloroplasts among the transformed protoplasts was significantly reduced by BFA treatment, similar to OsPHR (Supplemental Fig. S3). Moreover, to investigate whether proline residues in the N-terminal region of PHR are involved in chloroplast translocation, a transient expression vector encoding mutated TaOsPHR or HvOsPHR was constructed by replacing the 7th or 9th proline residue with an alanine residue (P7A-TaOsPHR, P9A-TaOsPHR, P7A-HvOsPHR, or P9A-HvOsPHR). The results showed that in both TaOsPHR and HvOsPHR, the conversion of the 9th alanine was inhibited to a greater extent than that 7th alanine (Supplemental Fig. S4). These results suggested that other *Poaceae* plants also have a PHR chloroplast translocation via the ER-Golgi system similar to that of rice.

## Discussion

Several chloroplast-localized proteins without predicted transit peptides are transported to chloroplasts through the secretory pathway of the ER-Golgi system (Kleffmann et al., 2004; Friso et al., 2004; Asatsuma et al., 2005). Our results showed that OsPHR, which is transported and functions in the nucleus, mitochondria, and chloroplasts, is transported to chloroplasts via the ER-Golgi system because BFA treatment inhibited OsPHR translocation to chloroplasts. Proteins transported to chloroplasts via the ER-Golgi system are characterized by 1) possessing a vesicular signal peptide in the N-terminal region, and 2) an N-terminal region rich in hydrophobic amino acids (Villarejo et al., 2005; Villarejo et al., 2006). The N-terminal region of not only OsPHR but also TaPHR, HvPHR, and ZmPHR, where citrine fluorescence was detected in chloroplasts, contains a sequence that is predicted to be an ER signal peptide by TargetP analysis and is more hydrophobic than those of *Arabidopsis* and other plants where PHR is not transferred to chloroplasts (Figs. 5A and Supplemental Fig. S1). Furthermore, amino acids with low helical propensities, such as proline, in the N-terminal hydrophobic amino acid region have been reported to be important for N-terminal vesicle transport (Kida et al., 2000; Kanaji et al., 2000). The N-terminal hydrophobic region of OsPHR contains numerous proline residues, and when the 8th and 9th proline residues were converted to alanine with a high helical propensity, translocation to the chloroplasts was significantly suppressed (Fig. 3). Conversion of the 7th and 9th proline residues of TaPHR and HvPHR to alanine markedly suppressed chloroplast translocation, similar to OsPHR (Supplemental Fig. S4). These results indicate that proline residues in the N-terminal region of PHR are important amino acid residues for N-terminal vesicular transport.

Furthermore, we showed that the phosphorylation state of the 7th serine adjacent to the 8th and 9th proline residues is strongly involved in the chloroplast translocation of OsPHR (Fig. 4). Phosphorylation has been reported to not only cause changes in protein conformation and conformational dynamics (Johnson et al., 2009; Park et al., 2010; Kast et al., 2010) but also affect the localization of proteins in the cell (Kuriyan et al., 2007; Pooler rt al., 2010; Gelvin et al., 2010; Ewens et al., 2010; Muñoz-García et al., 2010) because amino acid phosphorylation changes the structure and physicochemical properties of the amino acid itself. Thus, changes in the structure and physicochemical properties of the N-terminal region owing to serine phosphorylation may inhibit its passage through the ER. In this experiment, 7th serine of OsPHR was converted to alanine in the pseudo-unphosphorylated state, and the serine was converted to aspartic acid or glutamic acid in the pseudo-phosphorylated form. Alanine is slightly more hydrophobic than serine, whereas aspartic and glutamic acids are highly hydrophilic. The resulting suppression of chloroplast translocation of the pseudo-phosphorylated form of OsPHR may be due to the increased hydrophilicity of the 7th amino acid, which inhibits the chloroplast translocation of OsPHR through the ER. According to the molecular hydrophobicity potential analysis, phosphorylation is predicted to cause protein surface rearrangements and a local decrease in hydrophobicity near the phosphorylation site (Polyansky et al., 2012). Thus, there is a possibility that phosphorylation of the 7th serine affects not only the polarity of the serine itself, but also the neighboring proline. The resulting increase in hydrophilicity may inhibit chloroplast translocation through the ER. However, it is not clear what changes in the structure and properties of OsPHR are caused by phosphorylation of the 7th serine residue. These are important future issues in elucidating the molecular mechanisms of vesicular transport systems.

Previous analyses have shown that most OsPHR are present in a phosphorylated state (Teranishi et al., 2008). This may indicate that PHR is phosphorylated immediately after synthesis to inhibit its transport from its unphosphorylated state through the ER to the chloroplast. Phosphorylated PHRs, whose transport to chloroplasts is inhibited by phosphorylation, are then transported to the nucleus or mitochondria using C-terminal nuclear or mitochondrial transport signal sequences (Takahashi et al., 2014), respectively. However, when PHR is transported to chloroplasts, it is important that phosphorylated serine is dephosphorylated to a non-phosphorylated state, which allows PHR to be rapidly transported to chloroplasts via the ER. Thus, we propose that the distribution of PHR in the nucleus, mitochondria, and chloroplasts is regulated by the phosphorylation state of serine at position 7 (Fig. 6).

**Figure 6.**
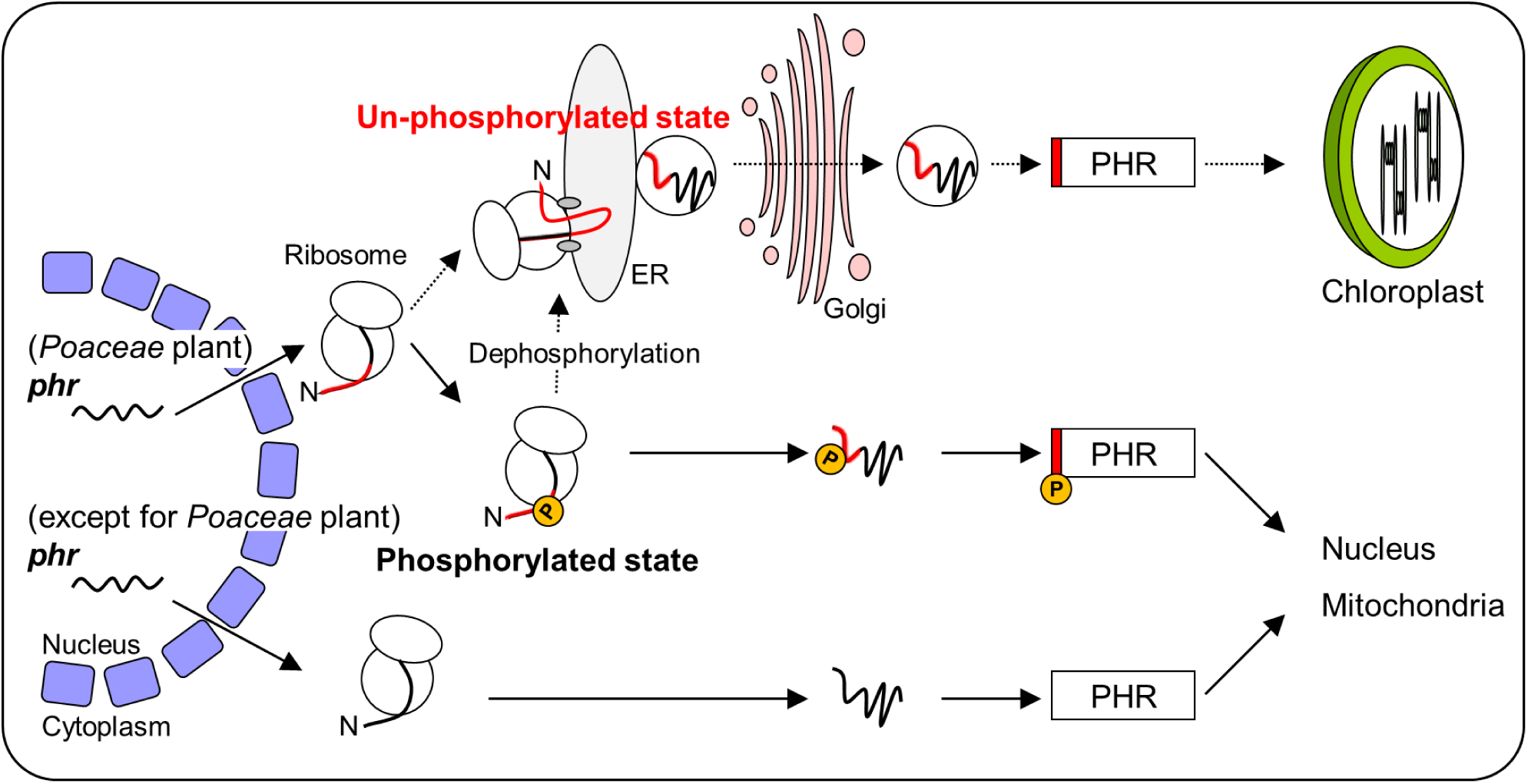
A speculative model of vesicle-mediated ER to Golgi to chloroplast PHR transport under the control of the sequence characteristics of N-terminal part of PHR. In *Poaceae* plants with proline-rich N-terminal sequences, the nuclear-encoded *phr*, in an unphosphorylated or dephosphorylated state, is transported in vesicle-mediated ER to Golgi then to chloroplast, according to the amino acid residues critical for N-terminal chloroplast translocation. On the other hand, PHR is not transported to the chloroplast when it is phosphorylated after translation initiation on the ribosome but is transported to the mitochondria and nucleus. Furthermore, in plants other than the *Poaceae* family, PHR is not transported to the chloroplast because it does not have the amino acid sequences important for translocation to the chloroplast on the N-terminal side and is transported to the nucleus and mitochondria according to the nuclear localization signal (NLS) and mitochondrial targeting signal (MTS) on the C-terminus. The N-terminal sequence enriched for proline residues is shown in red. Phosphorylated serine is shown by a yellow circle with the letter P.

No transfer to chloroplasts was observed in the PHRs of the plants used in this study, except for *Poaceae* plant PHRs. Several studies have examined the CPD repair activity in the chloroplasts and mitochondria of higher plants under visible light or dark conditions. Young *Arabidopsis* seedlings (5 days old) displayed no photorepair activity in their chloroplasts (Chen et al., 1996). Conversely, the leaves of *Arabidopsis* (14 days old) show blue light-dependent removal of CPDs from chloroplast DNA (Draper et al., 2000). However, it remains unclear whether this effect is mediated by CPD photolyase. Based on our results of 1) transient expression analysis of AtPHR using protoplasts prepared from rice or *Arabidopsis* seedlings (Figs. 5b and c), 2) analysis of the subcellular localization of AtPHR in transgenic *Arabidopsis* plants transformed with AtPHR (Figs. 5D and E), and 3) measurement of the activity of UVB-induced CPDs loss on chloroplast DNA (Fig. 5F), we concluded that AtPHR is not translocated or localized to chloroplasts in *Arabidopsis*. Kaiser et al. transiently transformed a GFP fusion of AtPHR (2 × CaMV 35S::AtPHR1::GFP) into green protoplasts from an *Arabidopsis* mesophyll cell culture to investigate the subcellular localization of AtPHR1. They demonstrated that the AtPHR1::GFP fusion protein was exclusively found in the nucleus and was not transported to the chloroplasts. These results support our findings.

In studies on the chloroplast localization of PHR in plants other than *Arabidopsis thaliana*, the number of CPDs in the chloroplast DNA of *Z. mays* (maize) leaves (Stapleton et al., 1997) was reduced in response to visible light. In contrast, no photoreactivation activity was detected in isolated *Spinacia oleracea* (spinach) chloroplasts (Hada et al., 2000). Although previous studies have suggested that plant species differ with respect to chloroplast localization of PHR, our results show that plant species differences in the chloroplast translocability of PHRs are due to differences in the presence or absence of sequences in the N-terminal region, which are important for translocation through the ER to the chloroplast.

## CONCLUSIONS

When transgenic rice plants, in which PHR do not function in chloroplasts, were grown under UV-B-supplemented conditions, leaf browning, a typical UV-B-induced chloroplast damage, was observed, which apparently caused growth defects (Fig. 1C). Therefore, it is necessary for PHR to function in the chloroplasts of rice plants to grow in the presence of UV-B, and rice plants may have acquired the ability to repair CPD in chloroplast DNA by transferring PHR to chloroplasts on their own during evolution. However, when *Arabidopsis* was grown under UV-B-added conditions, no browning of the leaves was observed, although the leaves showed UV-B damage that caused them to turn white (Dündar et al., 2020; Izumi et al., 2017; Otake et al., 2021). Thus, the UV-B damage phenotype is not necessarily uniform. These phenotypic differences indicate that the mechanisms of defence and adaptation to UV-B differ among plant species. For example, in *Arabidopsis*, whole chloroplasts damaged by UV-B irradiation are actively removed into vacuoles by chloroplast-targeted autophagy, also known as chlorophagy. Autophagy-defective *atg* mutants exhibit UV-B-sensitive phenotypes (Dündar et al., 2020; Izumi et al., 2017). However, such chlorophagic removal of impaired chloroplasts has not been observed in rice leaf cells with large chloroplasts, although rice also retains the chlorophagy mechanism, as plastids are removed by chlorophagy by vacuolar type H^+^-ATPase inhibitor concanamycin A treatment in the roots of non-green tissues in rice (Izumi et al., 2015). Thus, in plants without PHR translocation to chloroplasts, such a mechanism of chlorophagy may be activated to reduce UV-B-induced chloroplast damage. These differences in UV-B defence and adaptation mechanisms may also be due to the light environment in which each plant grows. In *Poaceae* plants grown under intense sunlight, maintaining mechanisms to repair chloroplast DNA damage may be more important than in plants grown under low sunlight. Nevertheless, the differences in the chloroplast localization of PHRs will be an interesting challenge in the future, as it is expected to lead to the elucidation of new UV-B defense/adaptation strategy mechanisms for growth in various light environments.

## Materials and methods

### Generating the expression constructs

The cDNA sequences of OsPHR (accession no. AB096003) and AtPHR (accession no. AF053365), or LjPHR (accession No. XP_057449869) was inserted into the pENTR/D-TOPO vector (Thermo Fisher Scientific, Waltham, MA, USA). To generate truncated or mutated forms of OsPHR, site-directed mutagenesis by inverse PCR was performed using the KOD-Plus Mutagenesis Kit (TOYOBO, Osaka, Japan) with OsPHR-inserted pENTR/D-TOPO as a template, according to the manufacturer’s instructions. Nucleotide sequences encoding the N-terminal positions of various plants were synthesized. The amino acid positions and accession numbers for each plant are as follows: TaPHR (amino acids 1-52, AK330529), HvPHR (amino acids 1-52, AK372010), ZmPHR (amino acids 1-39, NP_001130580), GmPHR (amino acids 1-42, NP_001238710), AmPHR (amino acids 1-44, ERN13316), SmPHR (amino acids 1-74, EFJ26958), and CrPHR (amino acids 1-79, AAD39433). The synthesized sequences were ligated to an expression construct encoding a truncated form of OsPHR (Δ_1-51_OsPHR) using restriction enzyme cloning or InFusion cloning (InFusion HD Cloning kit, Takara Bio, Shiga, Japan). The modified PHR was cloned into the binary vector pMpGWB106 (Ishizaki et al., 2015) via gateway cloning using LR Clonase II (Thermo Fisher Scientific). The expression constructs were driven by the CaMV 35S promoter with citrine fused to the C-terminus of the inserted sequence. The primers used for all plasmid constructs are listed in Supplemental Table 1.

### Production of transgenic plants

To produce transgenic rice plants, each prepared construct was transformed into the Agrobacterium strain EHA101. Transformation was performed as described previously (Hidema et al., 2007; Teranishi et al., 2004) using OsPHR gene-targeting (GT) rice plants generated as follows. For making the OsPHR GT rice, a 4.4 kb fragment containing the first exon from the promoter sequence and a 4.4 kb fragment starting from the first intron were amplified by PCR using the primers shown in Supplemental Table S1 (Supplemental Fig. S5A). The amplified fragments were cloned into the binary vector, pKOD4 (Saika et al., 2015). The construct contained neomycin phosphotransferase II (*nptII*) as a positive selection marker and the *diphtheria toxin A subunit* gene (DT-A) as a negative selection marker. The construct was then transformed into the Agrobacterium strain EHA105. Secondary calli induced from mature seeds of the rice cultivar Nipponbare (*Oryza sativa* L. ssp. *japonica*) were used for transformation (Teranishi et al., 2004; Saika et al., 2015). The transformed calli were selected by using G418 (35 μg mL^-1^). The plants were regenerated and then self-fertilized for two generations to obtain T_2_ plants. To confirm the gene targeting, PCR was performed using the primers listed in Supplemental Table S1 flanking the *nptII* insertion region (Supplemental Fig. S5B). Growth under supplementary UV-B radiation was tested to confirm the presence of GT plants (Supplemental Fig. S5C). Three independent lines were obtained using GT-c as the host in this experiment.

To produce transgenic *Arabidopsis*, the expression constructs were traed into the Agrobacterium strain GV3101. Agrobacterium was infected with the AtPHR-deficient mutant, *uvr2-1*, using the floral dipping method. Transgenic seeds were selected by using hygromycin B (50 μg mL^-1^). The plants were self-fertilized for two generations to obtain T_2_ plants.

### Plant materials and growth conditions

Transgenic rice plants were grown and treated with UV-B radiation as follows: seeds of each plant, soaked in water at 30 °C for 2 d, were sown in pots (15 cm wide × 6 cm deep × 10 cm high) containing fertilized soil in a large growth cabinet (Tabai Espec Ltd., Osaka, Japan), with a 12 h/12 h photoperiod and temperatures at 28/20 °C, as described previously (Mmbando et al., 2020). Photosynthetically active radiation (PAR) was recorded using a data logger (LI-1000; Li-Cor Inc., Lincoln, NE, U.S.A.) and an L1-190SA sensor (Li-Cor Inc.). PAR was adjusted to approximately 350 mol photons m^-2^ s^-1^ at the top of the plants. To assess UV-B sensitivity, plants were grown under visible radiation supplemented with UV-B radiation for 30 days using three UV-B bulbs (FL20SE; Toshiba, Tokyo, Japan) located above the plants. Plants receiving UV-B irradiation were subjected to the same photoperiod as the plants grown under visible radiation. Under UV-B bulbs, a UV29 glass filter (Toshiba Glass Co., Shizuoka, Japan) reduced 290 nm radiation by 50% (Kang et al., 1998). UV-B intensity was measured using a data logger (LI-1000) and an SD-104B sensor (Li-Cor Inc.). The UV-B intensity at the plant level is 1.0 W m^-2^. The spectral distribution was measured using a spectroradiometer (USR-45DA; Ushio Inc., Tokyo, Japan). Biologically effective UV-B radiation (14.7 kJ m^-2^ d^-1^) was calculated using the Caldwell plant action spectrum and normalized to unity at 300 nm (Caldwell et al., 1971).

To observed subcellular localization, the seeds were soaked in water at 28 °C for 5-7 days under visible white light (approximately 35 μmol photon m^-2^ s^-1^, as described previously (Takahashi et al., 2014).

*Arabidopsis* plants were grown in soil in chambers with a 16 h/8 h photoperiod and temperatures at 23 °C, under white fluorescent lamps (140 μmol m^−2^ s^−1^), as described previously^10^. Fourteen-day-old seedlings were used in all experiments.

### Preparation of protoplast from rice and Arabidopsis plant

Rice and *Arabidopsis* protoplasts were isolated as described by Kuroha et al., 2018 with minor modifications. Rice (*O. sativa L. Nipponbare*) plants were grown in plastic boxes with 1/2 Murashige and Skoog medium (Wako, Osaka, Japan) and 0.35% gellan gum (Wako) under white light condition at 30 °C for 10 days. *Arabidopsis* (*Arabidopsis thaliana Columbia-0*) plants were grown in pots containing fertilized soil under white light condition at 23 °C for 21 days. Leaf sheaths were used to isolate rice protoplasts, and leaves were used to isolate *Arabidopsis* protoplasts. After enzymatic digestion of each strip, the protoplasts were released by filtering through a 57 µm nylon mesh, followed by adjust approximately 4×10^6^ cells mL^-1^ with MMG solution (0.4 M Mannitol, 15 mM MgCl_2_ and 4 mM MES at pH 5.7) for PEG-mediated transformation.

### Protoplast transformation

Transformation was performed as described by Kuroha et al., 2018 with minor modifications. One hundred μL of isolated rice protoplasts were transferred into 1.5 mL microfuge tube including 3 μg of plasmid DNA and *Arabidopsis* protoplasts were transferred into 1.5 mL microfuge tube including 15 μg of plasmid DNA. After 10 min incubation on ice, 100 μL of polyethylene glycol (PEG) solution (40% PEG 4000, 0.2 M mannitol and 0.1 M CaCl_2_) was added, followed by incubation at room temperature for 30 min in the dark. After the incubation, 750 μL of W5 solution (154 mM NaCl, 125 mM CaCl_2_, 5 mM KCl, and 2 mM MES at pH 5.7) was added, mixed, and incubated under darkness at 25 °C overnight.

### Brefeldin A treatment

Brefeldin A (BFA) treatment was performed as described by Villarejo et al. (2005) and Isayenkov et al. (2011) with minor modifications. Stock solution of BFA (Sigma-Aldrich, Burlington, MA, USA) was prepared at 36 mM by dissolving BFA in DMSO. Protoplasts were treated with BFA (final concentration of 180 μM) by adding a stock solution 1 h after transformation. In this experiment, 0.1% DMSO was used as a control. After incubation under darkness overnight at 22 °C, the signal of Citrine-fluorescence in the BFA-treated or untreated protoplasts were observed using the laser-scanning confocal microscopy.

### Laser-scanning confocal microscopy imaging

Thin layers were peeled from the young leaves of the transgenic rice plants using needles and forceps. To detect mitochondria, the root was incubated with H_2_O containing 0.5 µM MitoTracker Red (Thermo Fisher Scientific) for 15 min at 25 °C, and then washed twice with H_2_O. Fluorescence was observed using confocal laser scanning microscopy (LSCM) performed using a C-apochromat LD63× water-immersion objective (numerical aperture = 1.15; LSM800, Carl Zeiss, Oberkochen, Germany). Citrine was excited using a 488 nm-laser, chlorophyll autofluorescence was excited using a 640 nm-laser, and MitoTracker Red were excited using a 561 nm-laser.

### Southern blot analysis

Southern blot analysis of organelle-specific DNA repair was based on a quantitative comparison of the intensity of T4 endonuclease V-treated and untreated DNA (Chen et al., 1996; Hanawalt et al., 1989; Stapleton et al., 1997). T4 endonuclease V cleaves single DNA strands at the CPD sites. Twelve-day-old L*er* seedlings grown on agar medium were irradiated for 15 min at a distance of 15 cm from the germicidal lamp (GL-15, Toshiba Electronic Co., Tokyo, Japan). The seedlings were then exposed to blue irradiation from blue fluorescent tubes for 10, 18, and 24h (FL20S B-F; National Co., Osaka, Japan), or were immediately placed in a light-tight box for 24h. The irradiation intensity was adjusted to approximately 60 µmol m^-2^ s^-1^. After then, the seedlings were immediately harvested and frozen in liquid nitrogen, then stored at -80°C until analysis. About 50 seedlings were used for each condition. Genomic DNA was isolated from the shoots of irradiated or non-irradiated seedlings, as described previously (Teranishi et al., 2004). Genomic DNA was digested using *Bam*HI. The digested DNA was divided into two equal aliquots; one aliquot was treated with T4 endonuclease V and the other was left untreated. The samples were resolved by alkaline agarose gel electrophoresis, and the DNA was blotted onto a nylon membrane as described previously (Hidema et al., 2007).

To generate probes for Southern blot analysis, a nuclear-specific gene (*NRPB1*: AT4G35800), a mitochondrial DNA containing mitochondrial-specific gene (*COX1*: ATMG01360) and a chloroplast-specific gene (*PSBA*: ATCG00020) were amplified using gene-specific primers (Supplemental Table S1). The amplified fragments were subcloned into pGEM-T Easy Vector (Promega). The digested inserts were labeled with [α-^32^P] dCTP and hybridized individually onto nylon membranes (Hidema et al., 2007).

### Statistical analysis

All data with error bars are presented as the standard error of the mean. Tukey’s test was used to compare multiple samples, as indicated in the figure legends.

## Supplemental Data

**Supplemental Figure S1.** Multiple sequence alignment of deduced amino acid sequences of PHR in various plant species using the Clustal omega program.

**Supplemental Figure S2.** Overall view of each membrane in Fig.5f.

**Supplemental Figure S3.** Cellular localization of *Poaceae* species PHR.

**Supplemental Figure S4.** Proline residues are important for chloroplast transition of *Poaceae* species PHR.

**Supplemental Figure S5.** Construction of OsPHR gene-targeting (GT) rice plants.

**Supplemental Table S1.** Primer sequences used in this study.

## Acknowledgements

We thank Dr. Atsushi Higashitani and Dr. Shusei Sato (Tohoku University) for their valuable discussions.

## Author contributions

M.O. and J.H. conceived and designed the experiments. M.O. performed most of the experiments. M.T. prepared the transgenic rice plants and measured CPD. C.K. and M.H. assisted in the subcellular localization analysis. K.Y. designed the constructs; M.O. M.T. and J.H. wrote the manuscript.

